# Proteomic analysis reveals complex interactions between PERK/ISR and MAPK1 in translational control in a phenotypic toxin model of Parkinson’s disease expressing PINTAC peptides

**DOI:** 10.1101/2024.12.14.628501

**Authors:** Girish N Nallur

## Abstract

In a proteomics investigation of a Parkinson’s disease [PD] cell model, a panel of 7 proprietary PINTAC peptides deployed in an *in vitro* phenotypic screen in SH-SY5Y cells treated with 6-OHDA identified 2 shorter variants of PERK protein kinase. One of the PERK variants, which is SUMOylated, associates *in vitro* with phosphorylated EIF2 alpha and its ubiquitinated isoforms, which also involves another PERK substrate. PINTACs revealed the presence of multiple post-translational modifications [PTMs] of MAPK1 in presence of OHDA. A fragment derived from MAPK1 protein sequence physically associated with multiple proteins from SH-SY5Y cells *in vitro*, including some PERK substrates involved in translational control. Most associations were abolished by OHDA treatment, with concomitant appearance of new interactions of the MAPK1 peptide with RSK proteins. Some PINTACs promoted the potentiation of the function of RPS6 *via* RSK activation induced by OHDA. A model of the phenotypic effects of aggregator and degrader PINTAC peptides is presented. The significance of these findings for novel drug discovery in PD is discussed in the context of target discovery and therapeutic applications of PINTAC peptides or small molecule mimics.

## Introduction

The progressive loss of dopaminergic neurons in the substantia nigra pars compacta [SNpc] accompanied by the depletion of dopamine [DA] levels in the striatum are the hallmarks of Parkinson’s disease [PD] pathogenesis [Masini *et al*., 2021]. Unfortunately, however, the molecular events which lead up to the PD pathogenic state are poorly understood. In spite of a firm genetic basis for PD pathology and the availability of candidate targets therefrom [Day *et al*., 2021], several decades of research and clinical investigation with many candidate drugs have had only limited success. Consequently, the true target(s) in PD may be still at large, elusive due to the lack of the right investigative tools and an absence of predictive screening methods.

Understanding the proteomic complexity of diseased states in extensive molecular detail, identifying damaged proteins which may be causative or contributive and generating useful biomarkers alongside the functions and biological contexts of dysregulated pathways in PD are desperately needed. Unfortunately, current molecular tools are utterly incapable of doing so, which is not the least of the confounding factors hampering progress. While it is invaluable to harness the key learnings from the past, these have typically led to shot-in-the-dark approaches based on limited knowledge, an unworthy strategy. Rather, finding the true target(s) in the context of the pathways wherein they function in PD diseased states increases the likelihood of future success.

An *in vitro* cellular toxin model of PD has been out of favor with some of the PD research community. However, it is important to realize that learning from such a model can be invaluable, particularly in view of the limitations associated with the availability of clinical samples [Shan *et al*., 2023]. Unlike the SH-SY5Y and 6-hydroxy dopamine [OHDA] cell-based toxin model wherein protein aggregation occurs within hours, potential neurotoxins may not achieve comparable concentrations or a sustained presence of protein aggregates in the brains of PD patients. In addition, the body possesses correctional measures whereby the biological impact of toxic insults may at least be partially alleviated or blunted. Therefore, it can be surmised that pathogenesis from neurotoxins of compounds [Shan *et al*., 2023] remains a likelihood in PD pathogenesis. Dismissing such mechanisms on the basis of the rapidity of attainment of pathogenic states in PD brains compared to *in vitro* systems may not have been prudent but may still have validity in screening.

This proteomic investigative report describes the application of a peptide tool set, designated as PINTACs, for phenotypic screening in the SH-SY5Y/OHDA model system. PINTACs [Nallur, 2022] are peptides that can activate a specific cellular enzyme to modify the target protein to which it binds. Ubiquitination of the PINTAC’s target(s) is the most common post-translational modification [PTM] resulting from PINTAC action. However, many additional, non-degradative modifications, including SUMOylation, ufmylation, neddylation, prenylation, nitrosylation, and ADP ribosylation, have been documented with mass spectrometry of PINTAC-modified targets [Nallur, unpublished observations].

Proximity-dependent action is a unique property of PINTACs whereby only those target molecules that co-localize with the enzyme machinery necessary for target modification under biological conditions achieve the modification. The target may, and frequently does, distribute to many additional sub-cellular locations. These are not altered/modified by PINTAC action unless they also have proximity to the same enzyme system at the second location. Therefore, PINTACs have compartmental selectivity not only to the target, but also to at least one component of the enzyme machinery at the address in the cell where the two interact. Consequently, PINTACs are precision tools for observing the pathways in which a target protein participates and documents target modification. As potential future clinical compositions, PINTACs can be anticipated to provide high levels of specificity and selectivity while eliminating off-location or off-target effects, which are some of the shortcomings of small molecule therapeutic agents. Further, protein phosphorylation is perhaps the only PTM which small molecules can address in a horizontal model of activation or inhibition. While some small molecules may be location specific, the vast majority lack specificity for a subcellular location or the ability to modify a clinical target with most other PTMs mentioned above [vertical model].

While the core technology of PINTACs was described previously, [Nallur, 2022] this report describes their application to phenotypic screening in a cellular toxin model of PD. The findings implicate PTM-containing enzyme isoforms and their substrates, rather than the native proteins themselves, to be relevant and involved in the aggregation of proteins and the modulation of signal transduction. Further, the findings suggest the potential of PINTAC peptides for therapeutic applications in PD or the design of candidate small molecules derived from the unique insights provided by PINTACs.

## Results

### Model system and strategy for targeting protein aggregation phenotypes with PINTACs

Initial studies showed that six hours of treatment of SH-SY5Y cells with the neurotoxin OHDA was sufficient to initiate the aggregated phenotype. The cells remained healthy and survived for at least an additional 48 hours in presence of OHDA. As shown in Figure 1, panel A, DNA constructs expressing PINTACs were introduced into SH-SY5Y cells by lipofection after six hours of pretreatment with OHDA and the incubation continued for an additional 48 hours in culture medium containing OHDA. The cells were harvested and cell extracts were prepared.

**Fig. 1.**
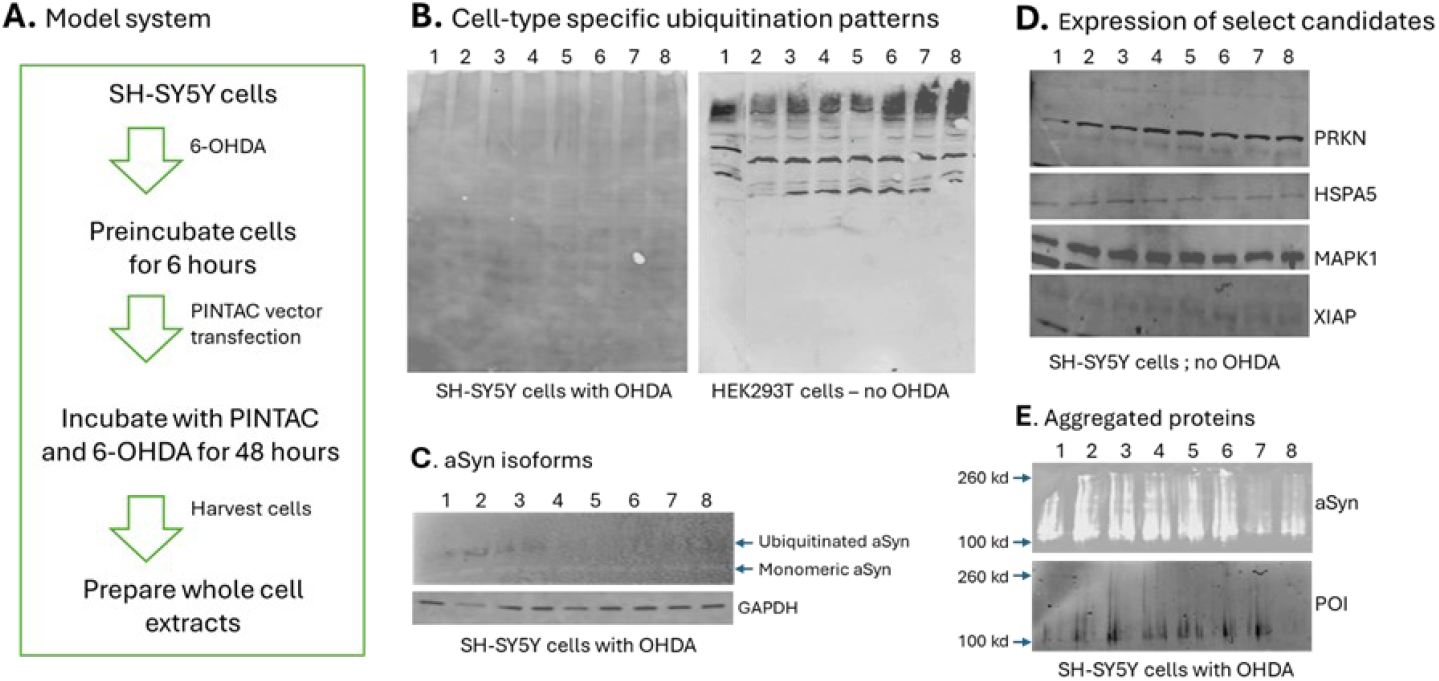
6-OHDA model system in SH-SY5Y cells and detection of protein modification in samples expressing PINTACs: PINTAC expression strategy in SH-SY5Y cells. Panel A – Schematic representation of the OHDA and PINTAC model system. Panel B – Western blot of total proteins with an antibody to ubiquitin. Panel C. Western blot with antibodies to αSyn and ubiquitin. Panel D. Western blot of select PD candidate proteins in SH-SY5Y cells without OHDA. Panel E – Western blot of SH-SY5Y proteins of high molecular weight showing aggregation in presence of OHDA. The numbers above each panel [throughout this report] represent samples expressing the following PINTACs: 1. Vector control; 2. PN001; 3. PN002; 4. PN003; 5. PN004; 6. PN008; 7. PN009; and 8. PN013.

Thus, the aim of this study was to observe the ability of PINTACs to overcome the toxic challenge from OHDA despite its continuous presence for the entire period of the study.

The expectation was that any PINTAC which targets a protein functioning in the aggregation pathway may exhibit a phenotypic effect in the OHDA model of PD.

### Ubiquitin modification by PINTACs follows a different pattern in SH-SY5Y cells when compared to HEK293T cells

The 8 PINTACs employed in this study gave rise to a robust smear of polyubiquitinated proteins ranging in size from 100 kDa to over 260 kDa in HEK293T cells [Figure 1, panel B, the polyubiquitinated fraction]. Mass spectrometry of polyubiquitinated proteins following treatment with several PINTACs identified targets unique to each PINTAC, as well as ubiquitin and several additional PTM tags known in the literature [not shown]. These studies indicate that treating cells with PINTACs can be considered as cell-based PTM assays. The substrate profiles of several E3 ligases, deubiquitinases, and other proteins of the ubiquitin proteasome system [UPS] were identified in this manner [not shown].

In contrast to HEK293T cells, the same PINTACs when expressed in SH-SY5Y cells yielded much smaller ubiquitinated proteins in the size range of 20 to 100 kDa, suggesting mono-ubiquitination [Panel B]. There was substantially less protein in the polyubiquitinated fraction with any of the PINTACs in SH-SY5Y cells when compared to HEK293T cells; moreover, the robustness of ubiquitination was substantially decreased. This may suggest that SH-SY5Y cells manage PTM mechanisms differently than HEK293T cells or that ubiquitination may be compromised in some manner in SH-SY5Y cells, particularly in the presence of OHDA. However, cell-type specific variations in the types of PTMs of specific proteins, including PD candidates, could not be ruled out [see below].

### PINTACs abolish ubiquitination of α-Syn and modulate its aggregation (up or down) in SH-SY5Y cells

Since the aggregation of α-Synuclein(α-Syn) is a marker of PD and other synucleopathies [Calabresi *et al*., 2023], it was important to determine if PINTACs can alter the PTM of α-Syn. As shown in Fig. 1, panel C, a basal level of monomeric α-Syn is expressed in SH-SY5Y cells regardless of the presence or absence of OHDA or PINTACs. α-Syn is ubiquitinated in OHDA-treated cells [lane 1] and in presence of PN001, PN002, and PN003. The remaining PINTACs abolished α-Syn ubiquitination.

Quantification of the signals from aggregated α-Syn [ag-α-Syn] from western blot images showed that PN001-004 increased the amount of ag-α-Syn by 2-to 4-fold when compared with OHDA alone [data not shown]. On the contrary, PN009 and PN013 reduced ag-α-Syn by 2x and 8x, respectively, within 48 hours of incubation [lanes 7 and 8]. This suggested that PN009 and PN013 may activate the UPS to degrade ag-α-Syn in an OHDA-specific manner since the same PINTACs showed no degradation of the small amounts of ag-α-Syn in SH-SY5Y cells [not shown]. Many additional proteins, including some PD candidates, are unaffected by PINTACs with or without OHDA, as shown in Fig. 1, panel D, suggesting a direct and specific impact on α-Syn, but not a generic response. These studies confirmed the ubiquitination of α-Syn proposed in the literature [Nonaka *et al*., 2005] and extended the findings with respect to the effect of PINTACs on α-Syn modification.

These observations suggested that PINTACs exert a phenotypic effect in SH-SY5Y cells targeted at specific proteins, raising the prospect that a deeper proteomic analysis of the OHDA system may elucidate the proteins and pathways involved in the response to PINTACs, particularly in the context of PD pathology. For this purpose, the samples were subjected to proteomic analysis in order to gain mechanistic insights into the pathways modified or the mechanism(s) of action of PINTACs with respect to α-Syn aggregation. In general, PN001-004 promote α-Syn aggregation, while PN009 and PN013 may be degraders.

### PINTACs selectively modify OHDA-induced integrated stress response [ISR] in SH-SY5Y cells concomitantly with activation of EIF2AK3 [protein kinase R-like ER kinase, PERK]

The integrated stress response [ISR] is a complex signaling mechanism employed by cells to overcome environmental stress or other pathological conditions [Costa-Mattioli & Walter, 2022]. ISR involves a system of at least four protein kinases, including PERK. When ISR is triggered, PERK is activated by phosphorylation which, in turn, phosphorylates its substrates [Ron, 2002] which leads to global inhibition of protein synthesis. EIF2S1 [EIF2 alpha] is a prime substrate of PERK, which attenuates protein synthesis after phosphorylation by PERK.

OHDA is a stressor which is known to trigger ISR in model systems as well as in the brains of PD patients [Lockshin & Calakos, 2024]. To determine if ISR is triggered in the OHDA treated cells, total proteins from OHDA- and PINTAC-treated samples were analyzed by western blotting using an antibody to PERK. As shown in Fig 2, panel A, two bands of apparent molecular weights 68 kDa and 48 kDa were detected, whose abundance varied in different samples, suggesting that PINTACs interfere with PERK expression and therefore modify the ISR response in OHDA-treated cells. However, a band corresponding to the size of native PERK [127 kDa] was not detectable in any sample. This is anticipated since, according to the manufacturer, the PERK antibody detects a band corresponding to over 140 kDa, which is likely a PTM-containing form of PERK. The 68 kDa isoform was also detected with an antibody to SUMO2/3, but not SUMO1. Accordingly, the 2 shorter isoforms were designated PERK-sf1-SUMO2/3 and PERK-sf2.

**Fig. 2.**
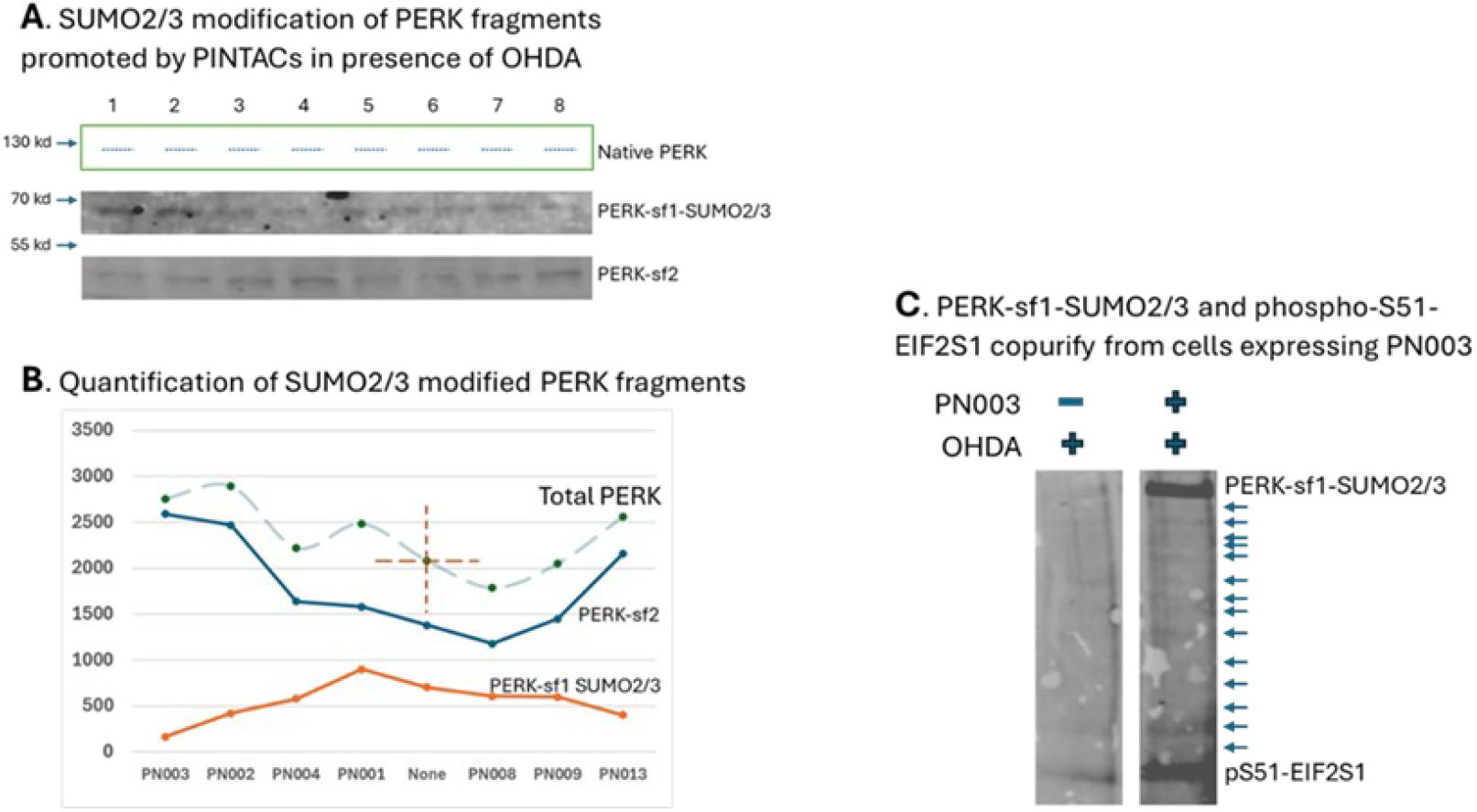
Evidence of PERK/ISR triggered by OHDA and modification of a shorter isoform of PERK with SUMO2/3. Panel A **–** Western blot of PINTAC and OHDA treated samples with antibodies to SUMO2/3 and PERK. A band of the expected size of native PERK was not detected with either antibodies [hence denoted by dashes]. However, a band of ^∼^68 kDa was detected with both antibodies suggesting a SUMOylated PERK isoform, and a second band of ^∼^48 kDa was detected with the PERK antibody alone. Panel B – Quantification of PERK isoforms and modeling of total PERK in the samples to differentiate the responses provided by different PINTACs for PERK modifications. Panel C – Stimulation of expression of PERK-sf1-SUMO2/3 by PN003 and its copurification with S51-phosphorylated EIF2S1 using an antibody to a protein of interest [POI] identified in this study. Arrows indicate putative isoforms of p-S51-EIF2S1 which were co-enriched in the assay.

PINTACs affected the expression of PERK-sf1-SUMO2/3 and PERK-sf2 isoforms. As shown in panel B, a marked increase in PERK-sf2 was observed with PN003 along with downregulation of PERK-sf1-SUMO2/3. PN013 also showed a similar distribution of the two PERK isoforms, but in contrast to PN003, which is an α-Syn aggregator [see Fig. 1 Panel E], PN013 provides a degrader phenotype. This is the first report of SUMOylation of PERK and the existence of shorter isoforms in OHDA-treated SH-SY5Y cells.

Curiously, the total PERK in the samples was nearly the same and comparable with OHDA alone, as shown by the dashed blue line and the red cross hatch, respectively. Total PERK values for aggregator PINTACs were deliberately plotted on the left of the cross hatch and the two which degrade aggregated α-Syn are on the right to emphasize that they have opposite phenotypic effects. However, the relative amounts of the PERK isoforms do not correlate with the pattern of PINTAC-induced changes to ag-α-Syn amounts [see Fig. 1], suggesting that other factors may better correlate molecular changes with aggregation. These observations prompted a deeper analysis of the proteomics of PERK in the OHDA model.

In an elegant series of experiments, Sassano *et al*. [2021] identified the substrates of PERK. Many of those proteins were detected by mass spectrometry of the polyubiquitinated fractions from PN008-, PN009-, and PN013-expressing HEK293T cells [not shown]. The same data also contained additional ISR enzymes, other stress responsive protein kinases which may aid or abet ISR enzymes, and components of the EIF2 complex. Therefore, this study views the observed changes in the OHDA model as stemming from ISR, and not limited to PERK activation.

The interplay and cross-talk between ubiquitination and phosphorylation to regulate signal transduction has been reviewed [Nguyen *et al*.; 2013], but a detailed description of the mechanism in any one system is lacking. The mass spectrometry data from this study showed that some PERK substrates may also serve as ubiquitination targets, being differentially modulated by PINTACs. To further explore the role of PN003 in the PERK/ISR process, the interactome of one PERK substrate suggested by Sassano *et al*. was enriched by ‘pull-down’ from the OHDA samples. The pulled-down proteins were analyzed by western blotting with antibodies to PERK and phospho-S51-EIF2S1 [phosphorylated EIF2 alpha]. As shown in Figure 2, panel C, PERK-sf1-SUMO2/3 and the phosphorylated isoform of EIF2S1, phospho-S51-EIF2 alpha, [Adomavicius *et al*., 2019] copurified with the pulled-down proteins from the PN003/OHDA sample, suggestive of a strong physical interaction with the PERK substrate identified by Sassano *et al*.. In contrast, relatively smaller [basal] amounts of these proteins copurified from the PN003/no OHDA, PN013/no OHDA or OHDA alone samples. These observations indicate the formation of a complex between the PERK substrate identified by Sassano with PERK-sf1-SUMO2/3 and/or phospho-S51-EIF2 alpha, the significance of which remains to be determined.

A highly notable aspect of this investigation is that multiple PTM-containing isoforms of pEIF2S1 were detected specifically in the PN003/OHDA sample. The bands present with an approximate spacing of ∼10 kDa and were also detected by an antibody to ubiquitin when the blot was reprobed [data not shown]. These bands were absent from cells treated with OHDA alone or other PINTAC samples. Taken together, the observations suggest a physical association of the PERK substrate identified by Sassano with the PERK and p-EIF2S1 isoforms. In control experiments, the purified PN003 peptide did not associate directly with p-EIF2S1, its isoforms, or PERK. Therefore, it is unlikely they are artefacts of the model system or an aberration of PN003 function. Since PINTACs exert their effects by a proximity-based mechanism, and may magnify or arrest target-dependent events, it is anticipated that p-EIF2S1 isoforms exist in biological systems. Importantly, PN013 abolished all interactions between the PERK substrate and PER or EIF2S1 [and its isoforms]. This suggest not only that PN003 and PN013 are related in function, but also that these complexes can be modulated for therapeutic benefit with small molecules or peptides. Since PN003 is an α-Syn aggregator, it appears that the presence of the complexes may promote aggregation. Further analysis in PD samples is needed in order to detect their existence and to elucidate their functions in more detail.

### PINTACs detect PTMs of MAPK1 (Erk2) in presence of OHDA

Mitogen-activated protein kinases (MAP kinases) respond to environmental stressors and are potent effectors of neuronal cell death [Harrison *et al*., 2004]. OHDA induces phosphorylation of c-Jun N-terminal kinase [JNK] leading to apoptosis in neurons [Albert-Gasco *et al*., 2020] and is upregulated in post-mortem PD brains. However, the role of MAPK1 signaling in neuronal health is less clear. Therefore, the structure and interactions of the MAPK1 protein was investigated in the OHDA- and PINTAC-treated samples.

In the absence of OHDA, phosphorylated p42/44 MAP kinase protein [pMAPK1] was expressed as a single band accompanied by a slight variation of expression with different PINTACs when compared with cells treated with OHDA alone [Fig. 3, panel A]. In presence of OHDA, however, PINTAC expression had a dramatic effect on the pMAPK1 protein. A triplet of closely spaced bands appeared whose relative intensities varied in some PINTAC samples. The fastest migrating species, which was the predominant band in cells treated with OHDA [lane 1], was expressed 2-3x higher in PN001 and PN008 samples. Expression of the slowest migrating species was highest in PN004-, PN008-, and PN009-treated cells. A third band in the middle was only detected in cells exposed to PN002, PN004, and PN013. These sample-specific expressions and PTMs of pMAPK1 imply that PINTACs enable their detection by magnifying the response of protein modifying enzymes. The requirement of OHDA for alterations of pMAPK1 protein structures suggests the involvement of modifying enzymes whose functions themselves are altered by OHDA, as well as proximity of the PINTAC to the enzyme machinery to give rise to highly polymorphic forms of the protein. For example, the slowest migrating species in lanes 5-7, and the signal bleeding, may signify a PTM specific to these samples. Therefore, PINTACs display convergence or divergence of pathways with specific proteins and measure the impact of OHDA. Further experimentation is needed to decipher if the triplet of pMAPK1 bands arises due to transcription, alternate splicing levels or at the protein level by conjugation with other proteins.

**Fig. 3.**
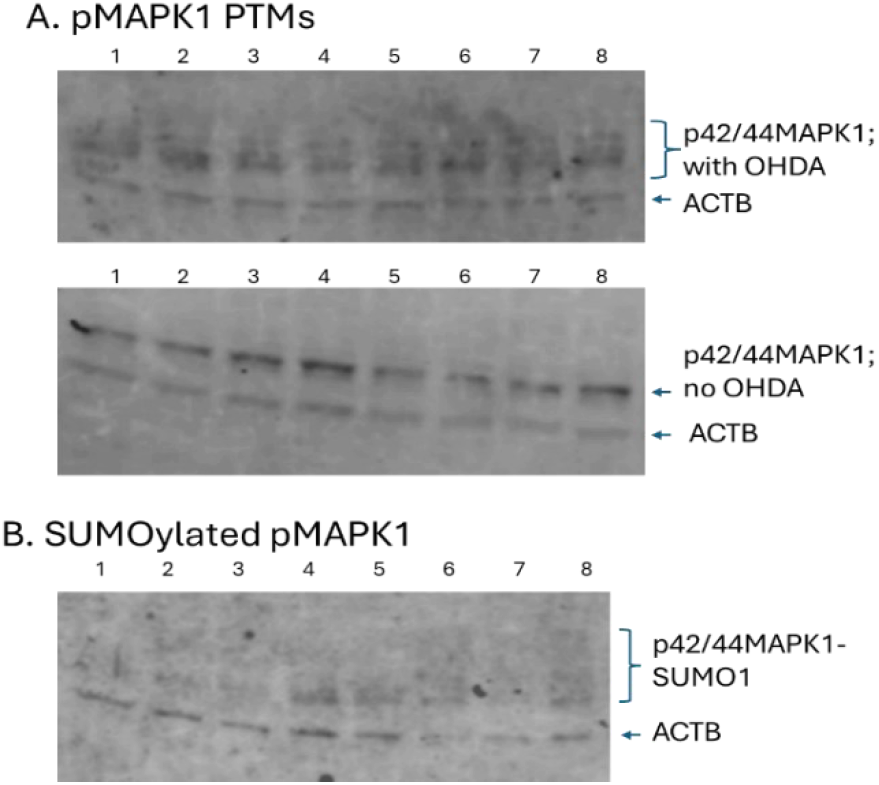
PTMs of p42/44-MAPK1. The indicated samples were detected with: A. an antibody to p42/44-MAPK1; B. SUMO1 antibody. Flower brackets indicate the multiple PTM containing forms of pMAPK1.

To further understand the effects of OHDA on the expression of pMAPK1 variants in SH-SY5Y cells, the samples were analyzed for the presence of PTMs of pMAPK1. As shown in Fig. 3, panel B, SUMO1 modification of the pMAPK1 variants was detected with some PINTACs. The fastest migrating variant of pMAPK1 was extensively SUMOylated in response to PN003, PN004, PN008, and PN013, and to a lesser extent by PN009. The ^∼^10 kDa increment in size by tagging with SUMO1 is consistent with the spacing of bands in the pMAPK1 triplet. Further, there were marked differences in the pattern of SUMOylated pMAPK1 products between α-Syn aggregator and degrader PINTACs. While only the fastest migrating species of pMAPK1 was SUMOylated by α-Syn aggregator PINTACs [PN003 & PN004], the α-Syn degrader PINTACs [PN009 & PN013] expressed a much broader smear of SUMOylation of all 3 pMAPK1 variants. These findings may suggest differences in functions or sub-cellular localization of PTM containing pMAPK1. The SUMO2/3 antibody did not bind to any of the MAPK1 variants.

The smear of SUMOylated bands observed with PN009 and PN013 suggest that there may be yet other modifications of pMAPK1-SUMO1 promoted by the α-Syn degrader PINTACs. However, SUMOylation alone does not explain the fuzzy bands which are of much smaller increments in size. Therefore, it is hypothesized that pMAPK1 in presence of PN009 and PN013 may involve additional modifications with smaller tags. Such tags include acetyl-, nitrosyl-, prenyl-, lipid conjugation, glycosyl-, or ADP-ribosyl whose tagging with pMAPK1 may better explain the size increments in the smear. If this type of protein maturation is confirmed *in vivo*, it may help to identify the primary pMAPK1 isoform as well as to explain the mechanism of actions of α-Syn aggregator and degrader PINTACs.

### Identification of a putative protein interactions site in MAPK1 adjacent to the RASopathy mutations

To begin to associate functional significance to the PTMs of pMAPK1, this study took advantage of mass spectrometry data from cells treated with PINTACs and the differences between HEK293T and SH-SY5Y cells [as shown in Fig 1, panel B]. The polyubiquitinated protein fractions from PN008, PN009, and PN013 assays in HEK293T cells were sequenced by mass spectrometry. As expected, MAPK1 [represented by 50 peptides] and MAPK3 [Erk1; represented by 65 peptides] were identified in the mass spectrometric data. Comparison of the tryptic peptide sequences of MAPK1 and MAPK3 identified a peptidic region of ^∼^50 aa which: a) maps C-terminal to the substrate binding site in MAPK1, beyond L241; b) is most divergent between the highly homologous MAPK1 and MAPK3 protein sequences; c) showed differences in putative PTM sites from PINTAC data; d) maps close to the RASopathy mutations site on its N-terminal site [Motta et al, 2020], and, e) contains 4 ubiquitination sites with reference to the PhosphoSitePlus [v6.75] database.

The ^∼^50 aa peptide derived from MAPK1, termed MK01-pep, helped to further analyze the PINTAC-induced PTMs in the OHDA cell model. This peptide was expressed in a human B cell line, DOHH2, purified, and used as a probe to identify associating proteins by far-western analysis. It is not known which PTMs, if any, the MK01-pep contains.

As shown in Fig. 4, a ladder pattern of 15-20 proteins [indicated by arrows, prominent bands are labeled] ranging in size from 9 kDa to over 130 kDa associated with MK01-pep in the absence of OHDA. PINTAC-induced differences in the signal intensities of several proteins were also evident.

**Fig. 4.**
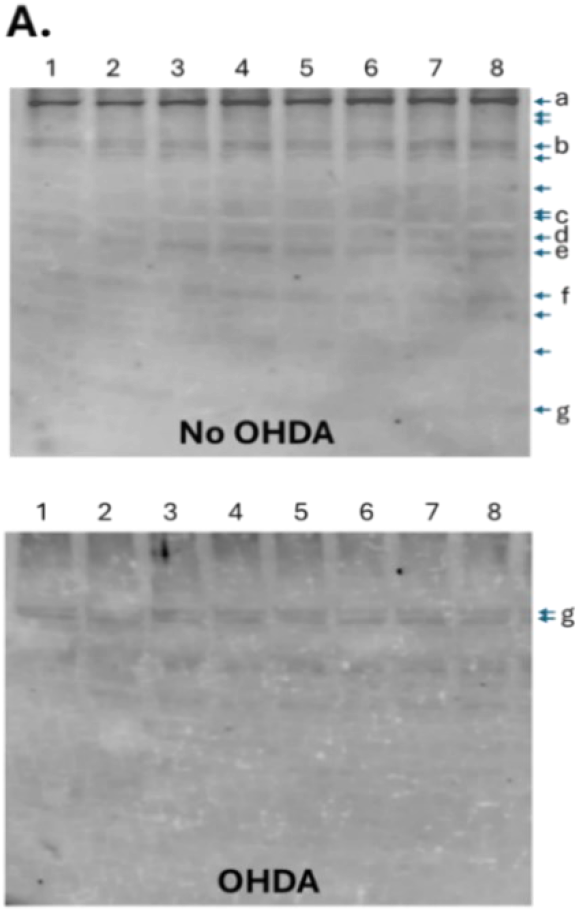
Physical interaction of MAPK1-pep in the OHDA model. Western blots of PINTAC treated samples with biotin-labeled MAPK1-pep. Arrows indicate the bands which associate with MAPK1-pep.

The profile of MK01-pep-associating proteins was dramatically altered with OHDA treatment [Fig 4. Panel A]. Many proteins no longer physically associated with MK01-pep, including the prominent bands ‘a’ and ‘b’. However, a new pattern of association emerged with OHDA treatment, with a doublet of bands marked ‘g’ [89-92 kDa in size] being the most prominent proteins. These findings confirmed that the MAPK1 peptide region constitutes a protein interactions motif, and that OHDA treatment modifies the expression or PTMs of many proteins resulting in loss of interactions.

### OHDA treatment of SH-SY5Y cells induces a cascade of protein kinase activation which is further modified by PINTACs

MAPK1 binds to and activates MAPK-activated protein kinases [MAPKAPs], including RSK1, RSK2, and RSK3, which, in turn, phosphorylate ribosomal protein S6 [RPS6] to modify protein translation. Tanoue *et al*. [2001] mapped the RSK1-interacting site in MAPK1 within a ‘docking groove’ in the protein with D316 and D319 involved in the interaction. Since the sequence of MK01-pep includes these amino acids near its C-terminus, it was anticipated to detect RSK1 in the OHDA model. Secondly, the apparent molecular weights of the protein bands marked ‘g’ in Fig. 4 are consistent with the sizes of RSK1/2/3.

To further explore the role of pMAPK1 in the OHDA-treated samples, the expression of total phosphorylated RSK1/2/3 was measured with an antibody and its physical associations with the biotin-tagged MK01-pep. As shown in Fig. 5, panel A, the RSK antibody detected both the bands marked ‘g’ in Fig. 4. There was a slight decrease of RSK expression in the PN001 and PN013 samples as compared with OHDA alone. Other PINTACs did not substantially alter the expression of RSKs.

**Fig. 5.**
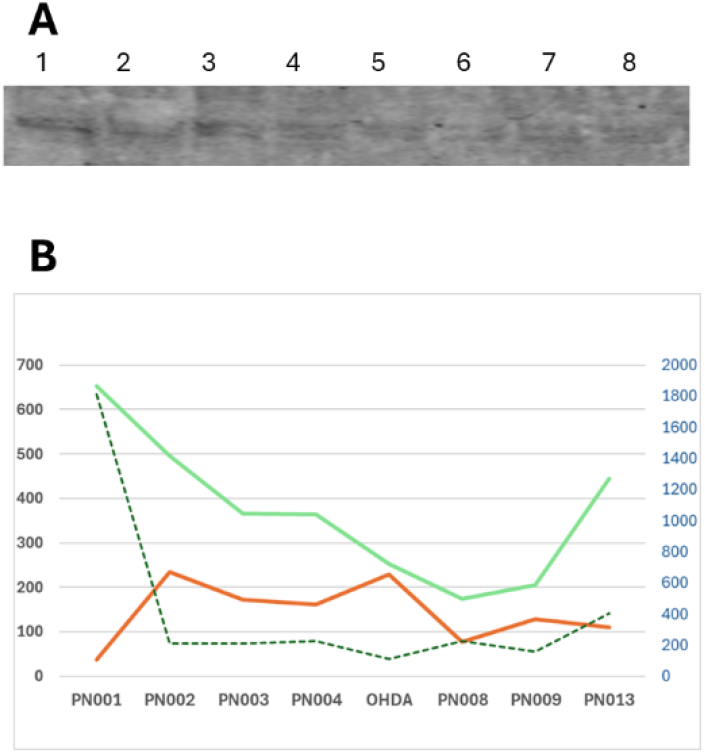
Comparison of the expression of RSKs and physical association with MK-01 pep. A. Western blot with an antibody to pRSK1/2/3. B. Quantitation of the signals from pRSK antibody [red line] and MK-01 pep binding to the upper band [green line]. The dotted line shows ratios of the two values the on the alternate axis.

On the other hand, the MK01-pep probe associated specifically only with the upper RSK band, with association strongest in the PN001- and PN013-treated samples.

When the signal intensities were plotted as a function of MK01-pep interaction normalized to unit RSK protein concentration, PN001 and PN013 exhibited the largest differences [Panel C, dotted line]. Note that PN001 inhibits PTMs of MAPK1 [see Fig. 3, OHDA samples, lane 2] while PN013 promotes it [lane 8]. This suggests that RSK may exist in different conformational states which one or more MAPK1 isoforms are able to distinguish and which PINTACs can modify.

Mass spectrometry of ubiquitinated proteins from PINTAC PN013-expressing HEK293T cells confirmed ubiquitination of RSK1 at K410 and K649 [data not shown]. However, additional putative PTM sites in RSK1 were also identified by mass spectrometry.

Purifying the RSK proteins in the ‘g’ doublet may help better understand their function as well as the consequences of the interaction with MAPK1. The MAPK1-RSK interaction may be a point of intervention for therapeutic applications in Parkinson’s disease. Further analysis of PN001 and PN013 in diseased samples and correlating differences with aggregated phenotypes may provide insights for development.

Some clues that the impacts of PN001 and PN013 on RSKs may not be equivalent came by observing the status of RPS6 in these samples. RSK1 phosphorylates RPS6 at all 6 serine residues, including Ser235/236. RPS6 is a critical regulator of ribosome biogenesis due to its role in translation of 5’ terminal oligopyrimidine [5’ TOP] tract containing mRNAs in a rapamycin-sensitive manner. 5’ TOP-containing mRNAs encode the translational machinery [Hagner *et al*., 2011]. The authors co-immunoprecipitated mRNAs containing 5’ TOP sequences with an RPS6 antibody, thereby demonstrating association of RPS6 with mRNAs.

In this study, total RPS was measured in samples expressing PINTACs. As shown in Fig. 4. Panel D, RPS6 was detected in all samples regardless of the presence or absence of OHDA, albeit with slight differences in expression. However, the bands migrated in gels at an apparent molecular weight of ^∼^55-60 kDa, suggesting that they may be dimers which are covalently conjugated under SDS-PAGE conditions. A band corresponding in size to the monomer RPS6 was detectable in PN013 and, to lesser extents, in PN008- and PN009-expressing samples.

A continuous smear of signals extending from the monomer to the dimer was observed in all three samples [Fig. 6.]. The monomer or the smear was not detected with other PINTAC expressing samples or OHDA alone. The RPS6 protein in the smear is not likely modified with ubiquitin or SUMO as are many other proteins in this study. The likelihood that they are RPS6-RNA conjugates is being investigated. If so, then PN013 potentiates RPS6 to overcome OHDA-mediated toxicity and recruit its mRNA substrates.

**Fig. 6.**
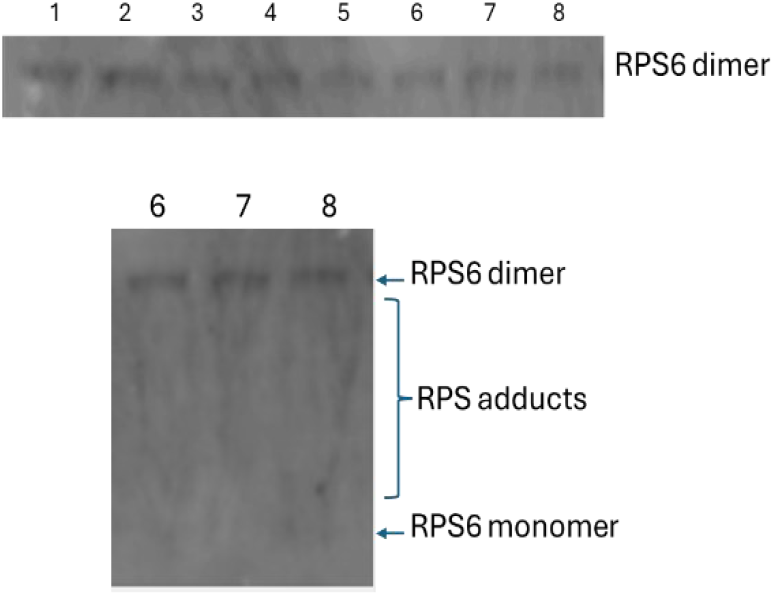
Detection of RPS6: Upper: Western blot showing the dimer of RPS6. Lower: The region of samples 6-8 showing the smear of signals between the indicated monomer and the dimer.

The unique ability of PINTACs to magnify, and sometimes arrest, proximity events may have caused the conjugates to accumulate which mat not occur in cells. If so, then the PN013-induced RPS6-RNA conjugates can be purified and studied further to enable drug discovery research. It is important to note that none of the α-Syn aggregator PINTACs, including PN001 which also stimulates RSKs to the same extent as PN013, exhibited the smear with the RPS6 monomer, suggesting that PINTACs modify RSK functions in different ways.

### MAPK1-pep may serve as a scaffold to coordinate the functions of protein complexes in translation, vesicular transport, redox control, and cytoskeletal organization

To further delineate the physical interactions of MAPK1-pep, the biotin-tagged peptide was used as a hook to pull down proteins from a whole cell extract of DOHH2 cells, a human B cell lymphoma cell line of the DLBCL type. Mass spectrometry of the pulled-down proteins yielded 85 proteins with a peptide count of at least 3, represented by 2,585 peptides. The data were queried for proteins with functions relevant to the OHDA PD model. Any protein with a peptide count of 5 or higher was arbitrarily considered to be a direct interactant with MAPK1-pep. Proteins represented by less than 5 peptides were only included in the analysis if the likelihood existed that they may able to form a complex with some other protein in the dataset which is represented by 5 or more peptides. The data identified 28 proteins involved in transcription and chromosome maintenance, including 9 histones and a histone binding protein. MAPK1 phosphorylates histones as part of its function in transcription and chromatin regulation [Suganuma and Workman, 2012]

However, MAPK1 has important roles to play in ER stress and protein folding and vesicular transport from ER to the Golgi. Therefore, the MAPK1-pep interacting proteins in the dataset are mentioned herein. The largest group of 13 proteins are involved in cytoskeletal organization [Table 1]. ACTB, MYH9, and TPM3 represented by 16, 19, and 16 peptides, respectively, are likely direct interactants with MAPK1-pep. Cofilin [CFL1], which is implicated in neurodegenerative diseases such as PD, Alzheimer’s and Huntington’s diseases was another key member of this group. ACTB is implicated in Baraitser-Winter syndrome and TPM3 in nemaline myopathy. Their roles in the OHDA model need to be investigated further. Moreover, ACTB and CFL1 are substrates of PERK according to Sassano *et al*. This study did not observe alterations of ACTB other than ubiquitination.

**Table 1.**
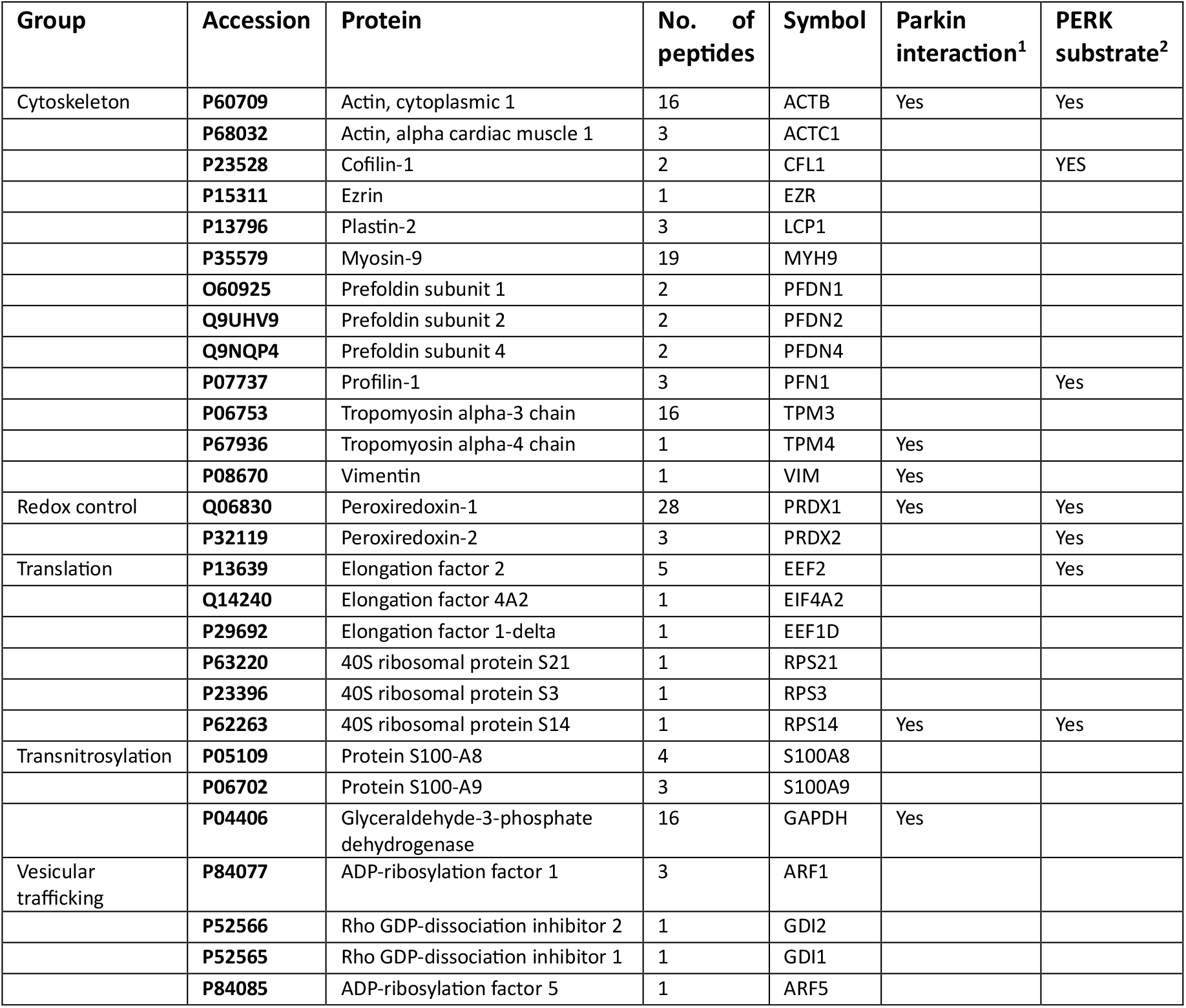
Proteins which physically interact with MAPK1-pep identified by mass spectrometry.

Translation factors and ribosomal proteins comprised the second largest group of 6 proteins. Elongation factor 2 [EEF2] is associated with spinocerebellar ataxia 26 [SCA26] but may also be involved in broader spectrum of NDDs. A variant of MAPK1 is the gene mutated in RASopathy and the mutations lie close to the coding region of MAPK1-pep. It remains to be determined if some of the interactants in this study could be involved in NDD as well. EEF2 may be a direct interactant of MAPK1; a band corresponding to the expected size of EEF2 was noted in the samples from this study, but not characterized.

Rare EIF4A2 variants cause intellectual disability, hypotonia, and epilepsy. By extension of the EIF2S1 interactions described earlier, RPS14, PRDX1, and ACTB have been shown to be substrates of PERK as well as Parkin. Therefore, it is intriguing that these proteins were physical associated with MAPK1, particularly in view of the number of different MAPK1 PTMs identified in this study.

Several S100 proteins copurified with the MAPK1-pep interactants. Notable candidates of interest in PD include S100A8 and S100A9. These proteins form a complexes with iNOS which is involved in MEK-ERK signal transduction [Jia *et al*., 2014]. The complex can direct S-nitrosylation of GAPDH, EZR, and VIM, which, except for iNOS, were identified among the mass spectrometry data.

Vesicular trafficking processes are disrupted in PD [Ebanks *et al*., 2020]. Therefore, it is intriguing to note that the MAPK1-pep pulled down ARF1/5 and GDI1/2. These proteins need to be analyzed in more detail in the OHDA- and PINTAC-treated samples.

An important point to note is that none of the proteins compiled in Table 1 are described in the literature as interactants of MAPK1. Perhaps the reason for this is that the protein interaction data were obtained with native proteins. As evident throughout this study, proteins containing PTMs frequently have a different interactome. These findings suggest that MAPK1 may act as a ‘scaffold’ to tether protein complexes to the cytoskeleton. Indeed, RSK1 has been suggested to play a role in actin reorganization after phosphorylation of Ser2152 by PAK1 [Vadlamudi *et al*., 2002]. In relation to this, mass spectrometry data from HEK293T cells identified PTM sites in PAK1 with PN008 and PN013. An important next step is to identify the isoform of MAPK1 which is involved in these pathway modifications since there may be many isoforms.

### A predictive model of aggregation based on the phenotypic effects of PINTACs

The major advantage of scale in biological investigations is the ability to draw insights and predict outcomes. Mass spectrometry of the proteomic samples collected after cell treatment with various PINTACs can greatly expand the data set. Aggregated proteins of high molecular weights were present in many of the assays presented in this report, which can yield much more information about the proteins therein. However, an attempt has been made to model the available data using a 13-point analysis using α-Syn aggregation as the basis. According to the model in Table 2, a physiological state of SH-SY5Y cells wherein a combination of attributes is attained is predicted to be most likely to prevent aggregation. PN003 and PN013 exhibited the highest phenotypic effects when compared to treatment with OHDA, but led to opposite outcomes based on the metric of α-Syn degradation. The PERK/pEIF2S1 observations presented in this report is the major difference between the impacts provided by the 2 PINTACs. This suggests that additional mechanisms exist in the EIF2 complex which either promote or eliminate the aggregated phenotype, based on the physiological status of cells. Further studies are needed to understand its role in protein aggregation in PD or other NDD.

**Table 2.**
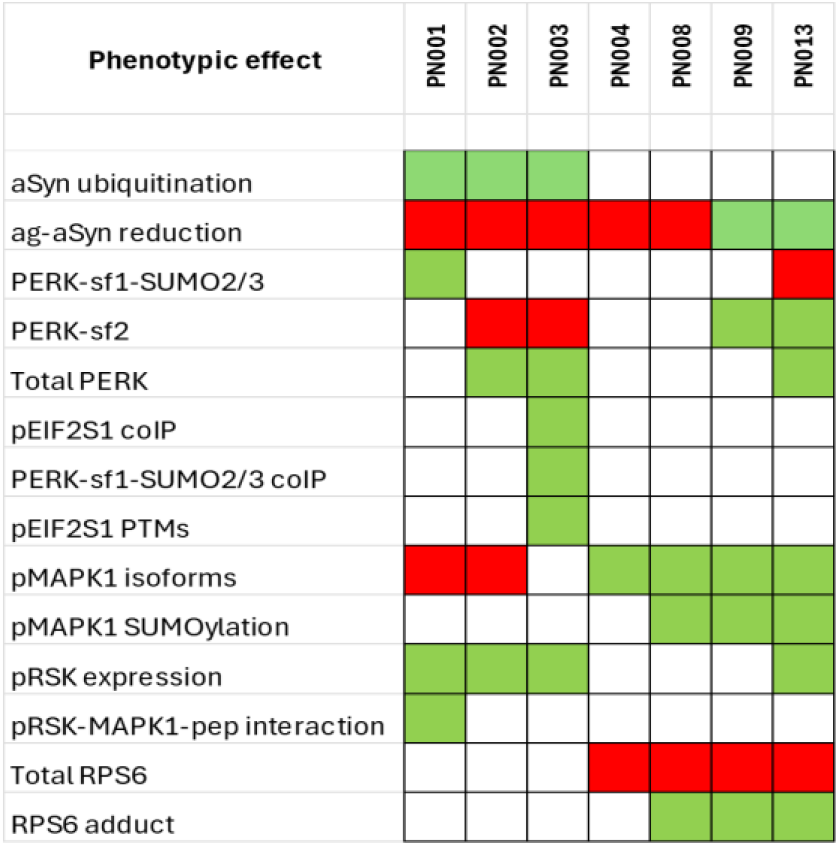
Heat map of the phenotypic effects of the PINTACs used in this study. Red-down-regulation; Green-upregulation.

## Conclusions

This is a first investigative report on the use of PINTAC peptides as a phenotypic screening tool. The 7 PINTACs used in this study were initially screened and characterized in HEK293T cells. Mass spectrometric analyses of the products generated by them included many proteins of interest in PD when compared with relevant literature. Therefore, it was compelling to apply them in the SH-SY5Y/OHDA PD *in vitro* cellular model used in this study.

The findings confirmed that PERK/ISR is activated, as suggested in the PD literature. However, the finding of 2 shorter isoforms of PERK, including SUMOylation of one of them, was unexpected. The strong *in vitro* association of SUMOylated PERK with its own phosphorylation target, pEIF2 alpha and its isoforms, was another surprise. It remains to be determined if this interaction constitutes product inhibition or if pEIF2 alpha has other substrates in addition to EIF2 beta. There can be many implications to therapeutics in PD and other NDDs if the same interactions are confirmed *in vivo* and in PD brains.

The finding that the PERK-pEIF2 alpha complexes are induced by the PINTAC PN003, which increases OHDA-induced α-Syn aggregation suggests that such complexes may need to be inhibited for therapeutic benefit in PD. The ability of PN003 to stably amplify the modified proteins can help develop assays for compound screening for novel PD therapeutics.

Many small molecules that activate or inhibit PERK have yielded highly ambiguous results. Our finding of multiple isoforms of the enzyme and its prime target, combined with the identification of a PINTAC which modulates the interaction, may offer insight into these ambiguous study results, as well as provide opportunities to develop assays for small molecule drug discovery.

Another serendipitous finding in this study is that there are multiple PTMs of MAPK1. It remains to be determined if the 3 isoforms of the protein observed with some PINTACs are alternately spliced forms. Interestingly, SUMO1 modification was also confirmed for some but the slowest migrating species likely have additional PTMs, such as ADP-ribosylation.

The physical association of MAPK1-pep with multiple proteins from SH-SY5Y cells and the abrogation of binding of most of them when exposed to OHDA suggests rampant changes to protein expression or structure by this neurotoxin. In future, pull-down experiments of the proteins associating with MAPK1-pep from multiple cell lines and comparison of mass spectrometry data will help to reconcile the impact of OHDA.

The specific binding of MAPK1-pep only to the RSK band with slower mobility, whilst the antibody detects both bands, suggests highly specific interactions between the enzymes. Nevertheless, PINTACs were able to distinctly elucidate differences in the binding intensity of the RSK-MAPK1-pep interaction, suggesting further heterogeneity of structures in the RSK band of lower mobility. Further analysis of this system employing the appropriate PINTACs will spur the identification of next generation inhibitors or activators.

In particular, the fuzzy smear of putative adducts associated with RPS6 specifically in the PN013-treated sample suggests that this study has begun to construct the full chain of control from OHDA through to the MAPK1-RSK-RPS6 axis and, very likely, down to the mRNAs recruited by RPS6 under different physiological conditions. This system can be invaluable for identifying the specificity of translatable mRNA in pathological conditions in PD and other NDD.

While every PINTAC used in this study offered a measure of unique proteomic insight, PN003 and PN013 exhibited the largest alterations toward the desired phenotypic effect. From the perspective of α-Syn, PN003 is an aggregator. Therefore, its impact on PERK and events surrounding the EIF2 alpha complex warrants further study. The finding that PN013 abolishes changes induced by PN003 suggests that the two are functionally related. α-Syn degraders have attracted special attention in the PD drug discovery space.

PN013 is an α-Syn degrader in spite of the highly challenging toxic environment induced by OHDA. Expansion of this study with a dose response to OHDA and incubating for longer periods may further delineate its role. If these studies are encouraging, PN013 has the potential to be developed as a therapeutic peptide if combined with a suitable delivery system.

Many additional peptide candidates were identified in this study which can be combined with PN013. As mentioned earlier, PINTACs have the unique property of affecting proximity to the target at a cellular address. Therefore, they hold considerable promise for developing drugs with absolute selectivity and little or no off-target effects.

In summary, the proprietary PINTACs deployed in the current proteomics studies and the biomarker roadmap developed in the phenotypic toxin cellular PD model offers a) opportunities to screen clinical samples from PD patients to validate our findings b) offers options to develop screening assays for SM drug discovery and lastly c) offers opportunities to optimize Pintac peptides (especially PN013) as therapeutics for suitable delivery to brain, and this will require medicinal chemistry skill deployment.

## Materials & Methods

DMEM (Cat. No. 11965-092), Penicillin-Streptomycin (Cat. No. 15140-122), Trypsin-EDTA (Cat. No. 25300-054) and Fetal bovine serum (Cat. No. A31604-02) were purchased from ThermoFisher Scientific. Lipofectamine2000 (Cat. No. 2352136) and Novex WedgeWell (Cat. No. XP00165) precast polyacrylamide gels were purchased from ThermoFisher Scientific. Rabbit antibodies against human Ubiquitin (Cat. No. 3933; Acc. No. P62987), PERK (Cat. No. 3192; Cell Signaling Technology), RSK1/2/3 (Cat. No. 9355; Cell Signaling Technology), pRPS6 (Cat. No. 4858; Cell Signaling Technology) and EIF2S1 (Cat. No. 3398; Cell Signaling Technology). Mouse monoclonal antibodies for MAPK1 (sc-514302) and aSyn (sc-12767) were purchased from Santa Cruz Biotechnology. 6-OHDA was purchased from SelleckChem (Cat. No. S5324).

### Cell culture, lipofection, whole cell protein preparation, and western analysis

SH-SY5Y cells were purchased from ATCC (Cet. No. HTB-11) and cultured according to manufacturer’s recommendations. Cells grown to mid-log phase were seed into culture dishes at 30% confluence and allowed to incubate for expansion up to 90% confluence. 6-OHDA was added to the cultured cells at a final concentration of 10 µM and preincubated for 6 hours prior to lipofection with the appropriate recombinant expression vectors harboring PINTAC sequences.

For lipofection: Cells were seeded in 100 mm tissue culture dishes at a density of 2.2 x 106 in 2 ml of growth medium (DMEM with 10% fetal bovine serum), and grown to a confluency of 90 per cent. Plasmid DNAs were introduced into the cells via transfection using Lipofectamine2000 according to the manufacturer’s protocol. Transfected cells grown for 72 hours at 37 °C under 5% CO2 without antibiotic selection.

For whole cell protein preparation, the lipofected plates were washed once with cold PBS. The adherent cells were resuspended in 400 ul of 1x Cell Lysis buffer (9803, Cell Signaling Technology) and incubated at room temperature for 15 min with occasional mixing. Lysates were cleared by centrifugation and the supernatants stored in aliquots at –20 °C.

Protein concentration of the cell extracts was determined using the Bradford-Lowry method (Quick Start™ Bradford Protein Assay Kit 1, Bio-Rad). For western blotting, whole cell proteins were separated on 12% SDS-polyacrylamide gels, transferred to nitrocellulose membranes and probed with the indicated antibodies according to western blotting protocols. Secondary antibodies tagged with infrared probes were purchased from Licor. Images were acquired using an Odyssey infrared scanner according to manufacturer’s recommended protocol.

### Sample preparation & mass spectrometry analysis of polyubiquitinated proteins

Whole cell extracts were fractionated alongside size markers on a 12% polyacrylamide gel containing SDS and stained with Coomassie Brilliant Blue R reagent in methanol and acetic acid reagent. After destaining, the bands above the 100 kd size were excised from gels and sent to mass spectrometry service provider for analysis. Mass spectrometry analysis was performed by the Biological Mass Spectrometry Facility of Robert Wood Johnson Medical School and Rutgers, The State University of New Jersey. Mascot software was used to identify proteins from LC-MS/MS data against the Swiss-Prot human database using a 1% false positive discovery rate (FDR).

### Mass spectrometry data analysis

The output from mass spectrometry (in Excel format) was sorted with respect to VB-009 (from high to low values) for the number of peptides identified against each protein. Proteins represented by at least 20 peptides, which is greater than 2 -fold as compared with the same protein in the pCMV6 control sample plus at least 2 other control PINTAC-treated cells were considered derived from proteins preferentially ubiquitinated in presence of VB-009.

## Acknowledgements

The author acknowledges strategic guidance from Drs. Dolatrai “Dinesh” Vyas, Nicholas A. Meanwell, and Ajay Bhargava and technical help from Dan Taranto.

## Notes

### Competing Interest Statement

The authors have declared no competing interest.

## References

1. Masini et al, 2021. Biomedicines, 9 598. A guide to the generation of 6-hydroxydopamine mouse model of Parkinson’s disease for the study of non-motor symptoms.

2. Day, JO and Mullin, S. 2021. Genes (Basel), 12 1006. The genetics of Parkinson’s disease and implications for clinical practice.

3. Shan L et al, 2023. npj Parkinson’s diseases, 9 #169. Towards improved screening of toxins for Parkinson’s risk.

4. Nallur, G, 2022. Research Square preprint. Induced polyubiquitination of proteins mediated by overexpression of a peptide: a novel tool for targeted protein degradation (TPD0 research. 10.21203/rs.3.rs-1547607/v1.)

5. Calabresi, P et al., 2023. Cell Death & Disease, 14 Article #176. Alpha-synuclein in Parkinson’s disease and other synucleopathies: from overt neurodegeneration back to early synaptic dysfunction.

6. Nonaka T, et al, 2005. Biochemistry, 44 361–368. Ubiquitination of α-Synuclein.

7. Costa-Mattioli, M & Walter, P, 2022. Science, 368 eaat5314. The integrated stress response: from mechanism to disease.

8. Ron, D, 2002. J. Clin. Invest. 110 1383–1388. Translational control in the endoplasmic reticulum stress response.

9. Lockshin ER & Calakos N, 2024. Current opinion in Neurobiology, 87 102886. The integrated stress response in brain diseases: a double-edged sword for proteostasis and synapses.

10. Sassano, ML et al., 2021. DOI: 10.1177/25152564211052392. Interactome analysis of the ER stress sensor PERK uncovers key components of ER-mitochondria contact sites and Ca2+ signalling.

11. Nguyen LK et al, 2013. Cell Commun. Signal. doi:10.1186/1478-811X-11-52. When ubiquitination meets phosphorylation: a systems biology perspective of EGFR/MAPK signalling.

12. Adomavacius, T et al., 2019. Nature Communications, 10 Article # 2136. The structural basis of translational control by eIF2 phosphorylation.

13. Harrison, JC et al., 2004. J. Biol. Chem. 279 2616–2622. Stress-specific activation mechanisms for the “cell integrity” MAPK pathway.

14. Albert-Gasco, H et al., 2020. Int. J. Mol. Sci., 21 4471. MAP/ERK signaling in developing cognitive and emotional function and its effect on pathophysiological and neurodegenerative processes.

15. Motta, M et al., 2020. Am. J. Hum. Genet. 107 499–513. Enhanced MAPK1 Function Causes a Neurodevelopmental Disorder within the RASopathy Clinical Spectrum.

16. Tanoue, T et al., 2001. The EMBO Journal, 20 466–479. Identification of a docking groove on ERK and p38 MAP kinases that regulates the specificity of docking interactions.

17. Suganuma, T & Workman, JL, 2012. Journal of Molecular Cell Biology, 4 348–350. MAP kinases and histone modifications.

18. Jia J et al, 2014. Cell, 16 623–634. Target-selective protein S-nitrosylation by sequence motif recognition.

19. Ebanks et al, 2020. Front. Neurosci., 13 1381. Vesicular dysfunction of Parkinson’s disease: Clues from genetic studies.

20. Vadlamudi, RK et al., 2002. Nat. Cell Biol. 4 681–690. Filamin is essential in actin cytoskeletal assembly mediated by p21-activated kinase 1.

